# Cortical Hierarchy Dynamically Organizes Large-Scale Neural Propagation

**DOI:** 10.64898/2026.08.29.747943

**Authors:** Xiaobo Liu, Bin Wan, Wenyue Liu, Runhao Lu, Sanwang Wang, Xihan Zhang, Qiuxuan Yu, Li Dong, Tengteng Fan

## Abstract

Flexible behaviour depends on the continuous coordination of sensory-driven and internally guided processing, yet whether the cortical hierarchy spanning lower-order sensory to higher-order association systems dynamically organizes large-scale cortical propagation over time remains unclear. Here we combined source-resolved magnetoencephalography with Riemannian cortical-flow modelling to derive hierarchy consistency, a moment-to-moment measure of the alignment between cortical propagation and the principal sensory-to-association functional gradient. We found that large-scale cortical propagation was dynamically organized by the cortical hierarchy. Hierarchy consistency exhibited a reproducible low-frequency periodic component that defined a characteristic timescale for the continuous updating of propagation direction. This dynamic organization was coordinated by a distributed cortical switchboard spanning the default-mode, salience, control and limbic systems, and was constrained by structural connectivity and network-control architecture. It flexibly adapted to behavioural demands, with hierarchy consistency increasing across both sensorimotor and working-memory states, while its characteristic periodicity shifted in a task-dependent manner. Moreover, hierarchy-related propagation dynamics were systematically reorganized across ageing and associated with higher-order cognitive function. Together, these findings establish the cortical hierarchy as a dynamic organizing principle that continuously shapes the direction and temporal evolution of large-scale cortical propagation to support adaptive behaviour.

## Introduction

The cerebral cortex is organized along a macroscale functional hierarchy that spans from primary sensory and motor regions to higher-order transmodal association cortex^1^. This hierarchical axis represents a fundamental organizing principle of cortical architecture, demarcating regions engaged in externally driven perception and action from those supporting integrative, abstract, and internally guided cognition^2,3^. Its expression is remarkably consistent across multiple levels of neural organization, encompassing patterns of functional connectivity, gradients of cortical microstructure and myelination, regional differences in receptor architecture, and systematic variation in cognitive specialization^4–9^. The sensory-to-association hierarchy thus provides a canonical spatial framework for mapping the large-scale functional organization of the human cerebral cortex.

Yet cortical organization is not solely a matter of spatial arrangement. Brain function is inherently dynamic: neural activity propagates across the cortical surface over milliseconds, generating transient spatiotemporal patterns that underlie perception, motor control, and higher cognitive function^3,10–13^. Whereas cortical gradient frameworks effectively describe the position of regions within the macroscale hierarchy, they are largely silent on how neural activity moves through this hierarchy over time^4^. A fundamental and unresolved question therefore remains: is moment-to-moment cortical propagation structured by the sensory-to-association hierarchy, or does the hierarchy principally reflect a static scaffold of functional organization that is largely decoupled from fast neural dynamics?

This question carries significant theoretical implications. Flexible, adaptive behaviour demands continuous and context-sensitive coordination between sensory-driven and internally guided processing^14–16^. Propagation aligned with the cortical hierarchy, from sensory toward association cortex, was interpreted as a putative bottom-up mode because its direction follows the canonical progression from externally driven sensory processing toward higher-order integration. Conversely, hierarchy-opposed propagation, from association toward sensory cortex, was interpreted as a putative top-down mode, consistent with feedback processes implicated in predictive modulation, internally guided cognition and cognitive control^14,16–18^. To support flexible cognition, the brain may therefore need to dynamically alternate between these two modes of activity flow^3,11,14^. Such alternation may depend on a distributed set of cortical regions that coordinate transitions between hierarchy-aligned and hierarchy-opposed propagation, thereby constituting a hierarchical switchboard for the dynamic reconfiguration of information flow across the cortical hierarchy. However, the temporal organization of such alternation, the cortical regions that coordinate it, and its functional relevance across the adult lifespan and for individual cognitive capacity remain largely unknown.

Here, we tested whether the sensory-to-association hierarchy provides a dynamic organizing axis for fast cortical propagation across intrinsic and task-evoked brain states^1^. Using source-resolved MEG and Riemannian cortical-flow modelling, we quantified the moment-to-moment alignment between cortical propagation and the principal functional gradient, termed hierarchy consistency. We then asked whether this hierarchical organization exhibits an intrinsic temporal structure, whether specific cortical regions coordinate its reconfiguration, and whether these dynamics are constrained by anatomical architecture. Finally, using complementary sensorimotor and working-memory datasets together with a large adult lifespan cohort, we examined how hierarchy-related propagation adapts to behavioural demands, changes across ageing and relates to individual cognitive ability.

## Results

### The cortical hierarchy provides a dynamic organizing axis for fast neural propagation

We first asked whether fast cortical propagation is systematically organized by the macroscale sensory-to-association hierarchy. Source-resolved magnetoencephalography (MEG) showed that local cortical activity spread directionally from high-activation centres to surrounding regions over millisecond timescales, forming structured propagation patterns across the cortical surface (**Fig. 1a**). To test whether these dynamics followed the macroscale functional hierarchy, we reconstructed instantaneous cortical-flow vector fields on the Riemannian cortical manifold and compared local propagation directions with the principal sensory-to-association functional gradient derived from resting-state functional magnetic resonance imaging (fMRI). Cortical propagation was not randomly distributed in directional space, but was systematically organized with respect to the sensory-to-association hierarchy. Propagation from sensory toward association cortex was defined as hierarchy-aligned, whereas propagation in the opposite direction was defined as hierarchy-opposed (**Fig. 1b**). Critically, this directional relationship was not static, but continuously reconfigured between the two propagation regimes, indicating that the sensory-to-association hierarchy provides a dynamic directional axis for fast cortical activity. We quantified this organization by measuring the instantaneous alignment between cortical-flow vectors and the local functional-gradient direction, termed hierarchy consistency. This measure captures the extent to which large-scale cortical propagation is organized along the sensory-to-association axis at each moment in time. Hierarchy consistency should therefore be interpreted as a time-resolved expression of hierarchical organization, rather than as a direct measure of synaptic communication or causal information transfer. We next asked whether this hierarchical organization exhibited an intrinsic temporal structure. Spectral parameterization of the hierarchy-consistency time series revealed a prominent low-frequency periodic component (**Fig. 1c**). Thus, alignment between cortical propagation and the functional hierarchy was not stationary, but was continuously reorganized between hierarchy-aligned and hierarchy-opposed regimes at a characteristic timescale. This low-frequency temporal structure was reproducible across the Cambridge Centre for Ageing and Neuroscience (Cam-CAN) and Open MEG Archive (OMEGA) cohorts, with robust spectral fits and clear inter-individual variability (**Supplementary Figs. 1 and 2**). Together, these results show that the sensory-to-association hierarchy not only defines the macroscale spatial organization of cortex, but also provides a dynamic organizing principle for fast neural propagation, with large-scale cortical activity continuously reconfigured between aligned and opposed propagation regimes over a reproducible low-frequency timescale.

**Figure 1.**
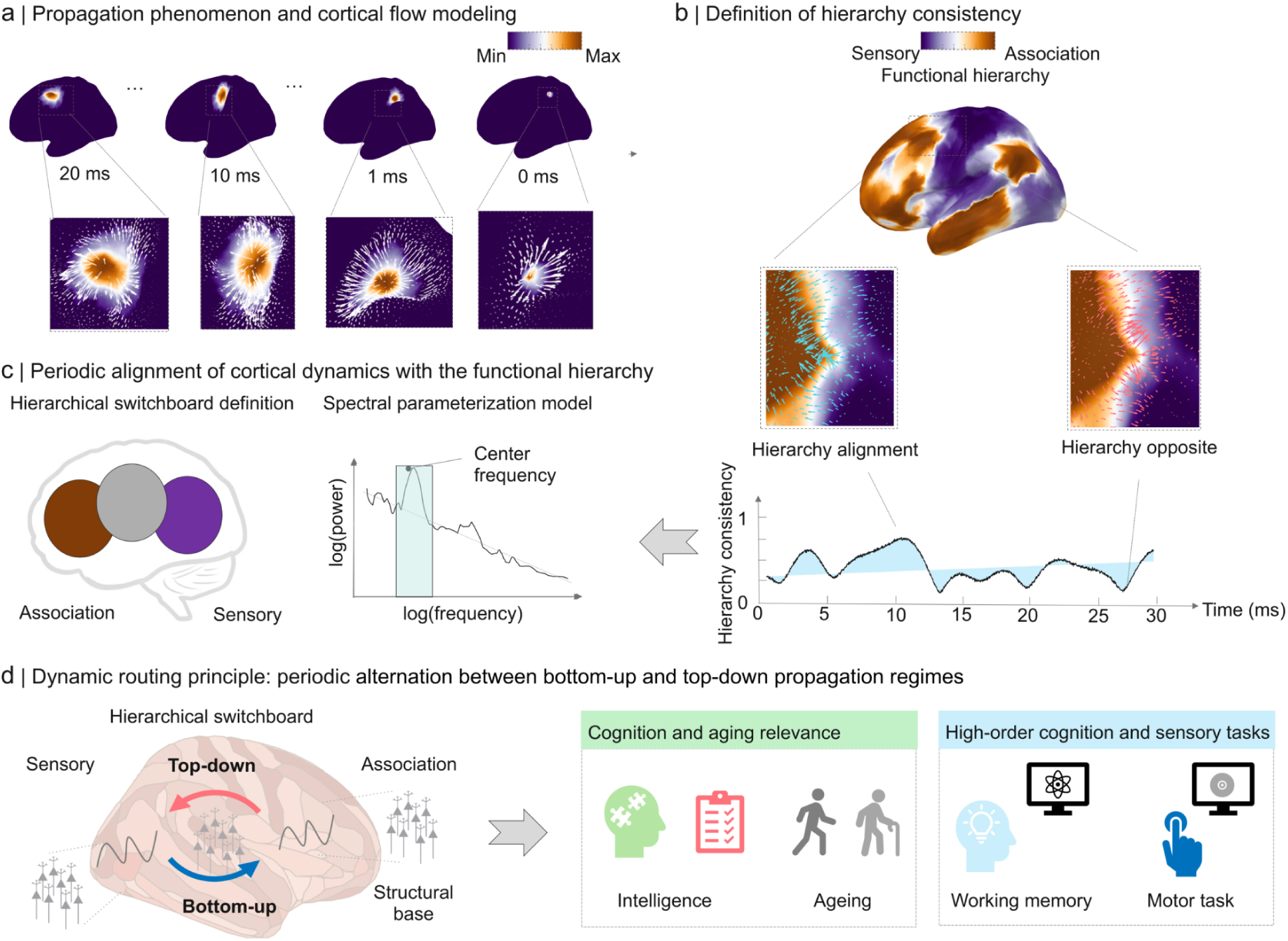
Cortical propagation, hierarchy consistency, and conceptual framework for hierarchy-related cortical dynamics. **a,** Propagation phenomenon and cortical-flow modelling. Selected cortical surface maps illustrate the spatiotemporal evolution of a local activation peak in source-resolved MEG. Insets show the corresponding cortical-flow vector fields, in which white arrows indicate the local direction and relative magnitude of propagation, and colour intensity from purple to orange denotes signal amplitude from minimum to maximum. These examples illustrate how local cortical activity gives rise to structured propagation patterns across the cortical surface. **b,** Definition of hierarchy consistency. The principal sensory-to-association functional gradient was used as a macroscale reference axis. At each cortical location, the instantaneous cortical-flow direction was compared with the local gradient direction. Propagation along the gradient was defined as hierarchy-aligned, whereas propagation against the gradient was defined as hierarchy-opposed. The resulting hierarchy-consistency time series quantifies moment-to-moment variation in the proportion of cortical propagation aligned with the functional hierarchy. **c,** Periodic alignment of cortical dynamics with the functional hierarchy. Left, schematic illustration of the hierarchical switchboard concept, showing a distributed cortical architecture linked to hierarchy-related propagation dynamics. Right, spectral parameterization of the hierarchy-consistency time series in log–log power–frequency space, used to separate periodic from aperiodic components and to estimate the centre frequency of the dominant low-frequency periodic mode. **d,** Conceptual summary of hierarchy-related cortical dynamics. The schematic illustrates periodic alternation between bottom-up (sensory-to-association) and top-down (association-to-sensory) propagation regimes, together with their proposed links to structural organization, ageing and cognition, and task-dependent modulation in both higher-order cognitive and sensorimotor contexts.

**Figure 2.**
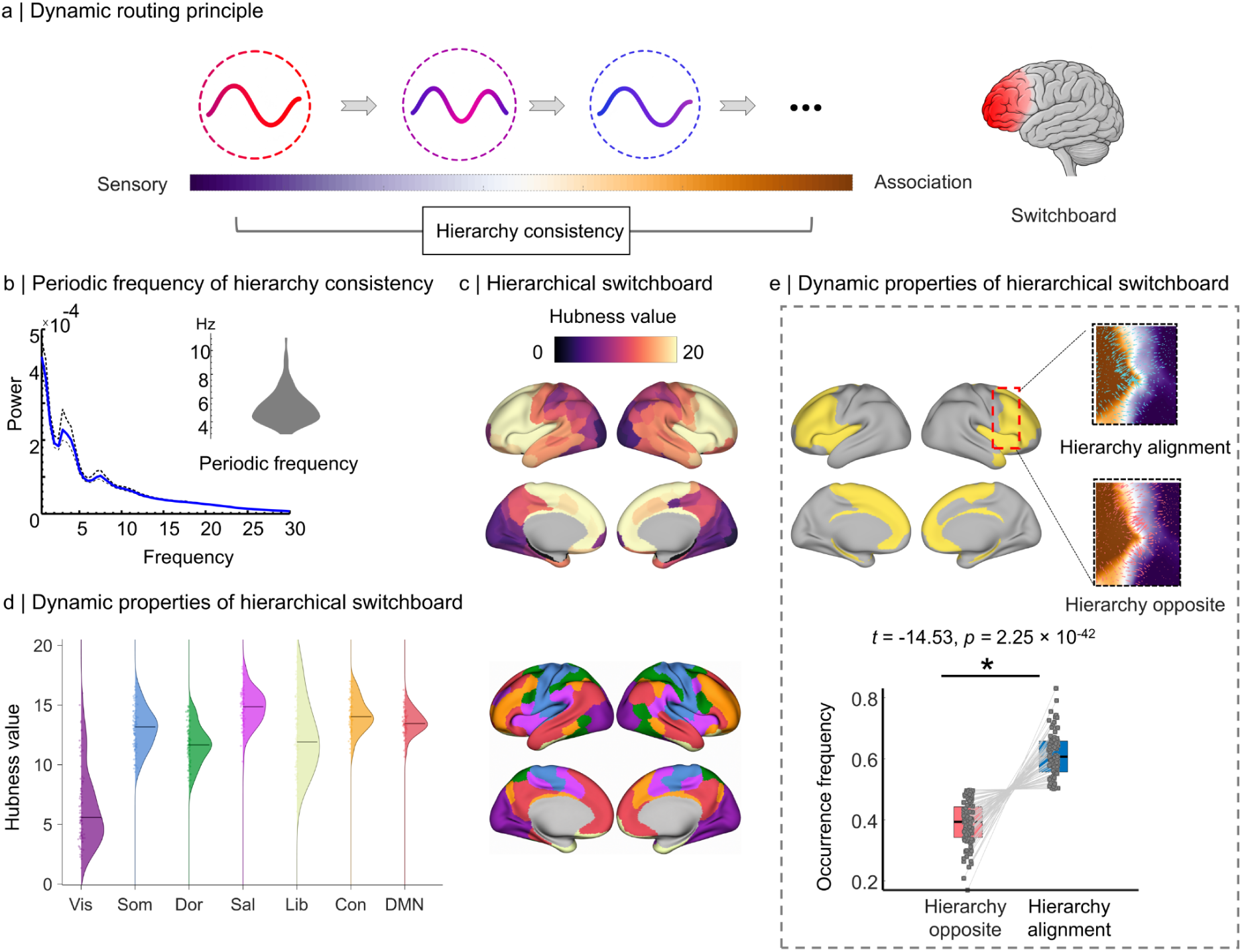
Periodic frequency of hierarchy consistency and spatial-dynamic properties of the hierarchical switchboard in OMEGA (N = 83). **a,** Schematic illustration of the dynamic routing principle. Moment-to-moment cortical propagation is evaluated relative to the sensory-to-association functional hierarchy, and its alignment defines hierarchy consistency. Sequential propagation states are shown along the sensory-to-association axis, with higher-order prefrontal association cortex highlighted as a major coordination site. **b,** Periodic frequency of hierarchy consistency. Power spectral analysis revealed a prominent low-frequency periodic component after accounting for the aperiodic background. The solid blue curve shows the group-mean spectrum, dashed black curves indicate the standard error, and the inset shows the distribution of individual centre frequencies (5.8 ± 1.4 Hz). **c,** Spatial distribution of hierarchical switchboard hubness. Cortical maps show parcel-wise hubness derived from the hierarchy-dynamics synchronization network. Higher hubness was preferentially distributed across higher-order association cortex, whereas lower hubness was more prominent in primary sensory regions. **d,** Network-level distribution of switchboard hubness. Hubness was higher in transmodal systems, including salience, frontoparietal control, limbic and default-mode networks, than in the visual network. Cortical maps show the seven canonical functional networks used for network-level characterization. **e,** Propagation-state expression within switchboard regions. Yellow masks indicate switchboard-related regions identified using *k*-core decomposition. Insets illustrate representative hierarchy-aligned and hierarchy-opposed propagation patterns. For the paired contrast hierarchy-opposed minus hierarchy-aligned occurrence, hierarchy-opposed states occurred significantly less frequently than hierarchy-aligned states (*t* = −14.53, *p* = 2.25 × 10⁻^42^).

### Hierarchy consistency exhibits a low-frequency periodic component and a cortical switchboard organization

Having established hierarchy consistency as a time-resolved measure of propagation alignment, we next examined its spectral structure (**Fig. 2a**). Power spectral analysis of the hierarchy-consistency time series revealed a low-frequency periodic component after adjustment for the aperiodic background (**Fig. 2b**). Individual-level centre frequencies were predominantly distributed within a low-frequency range (5.8 ± 1.4 Hz), indicating that the dynamic alignment between cortical propagation and the functional hierarchy is not merely a static property, but fluctuates rhythmically over time.

We next identified cortical regions whose hierarchy-related temporal profiles were centrally embedded within transitions between hierarchy-aligned and hierarchy-opposed propagation states. Using k-core decomposition of an inter-regional hierarchy-transition synchronization network(**Supplementary Fig. 3**), we derived parcel-wise hubness values indexing each region’s centrality in this transition-related network. The resulting hierarchical switchboard was spatially non-uniform: high-hubness regions were concentrated in higher-order and transmodal association cortex, whereas primary sensory regions exhibited comparatively lower hubness (**Fig. 2c**). Network-level analysis showed higher hubness in transmodal systems, including salience, frontoparietal control, limbic and default-mode networks, relative to primary visual cortex (**Fig. 2d**). These findings suggest that higher-order association systems are preferentially involved in coordinating transitions between propagation regimes along the sensory-to-association hierarchy.

To further characterize switchboard dynamics, we quantified the relative occurrence of hierarchy-aligned and hierarchy-opposed propagation states within switchboard-related regions (**Fig. 2e**). For the paired contrast hierarchy-opposed minus hierarchy-aligned occurrence, hierarchy-opposed states occurred significantly less frequently than hierarchy-aligned states (*t* = −14.53, *p* = 2.25 × 10⁻^42^), indicating that switchboard regions preferentially express propagation consistent with the macroscale functional hierarchy while retaining the capacity to enter the opposing regime. Together, these findings identify a spatially structured cortical switchboard associated with rhythmic alternation between hierarchy-aligned and hierarchy-opposed propagation states.

### Task engagement enhances hierarchy consistency and flexibly modulates its periodic dynamics

Finally, we examined whether hierarchy-consistency dynamics were flexibly modulated by different task demands. In the Cam-CAN sensorimotor task, hierarchy consistency was significantly higher during task than rest (*t* = 20.95, *p* = 1.65 × 10⁻⁶⁹), indicating stronger alignment of cortical propagation with the sensory-to-association hierarchy during externally driven sensorimotor processing. Periodic frequency was also significantly increased during task (*t* = 15.48, *p* = 1.43 × 10⁻^43^), indicating faster temporal reorganization of hierarchy-related propagation dynamics. In the HCP working-memory task, hierarchy consistency was likewise significantly higher during task than rest (*t* = 2.64, *p* = 0.0098), suggesting that higher-order cognitive engagement also strengthened hierarchy-consistent propagation. In contrast to the sensorimotor task, however, periodic frequency was significantly reduced during working memory (*t* = −2.68, *p* = 0.0086), indicating slower hierarchy-related dynamics during higher-order cognitive processing (**Fig. 3**). Together, these findings show that task engagement consistently enhances the alignment of cortical propagation with the functional hierarchy, while its characteristic temporal scale is flexibly tuned to task demands, accelerating during sensorimotor processing but slowing during working memory.

**Figure 3.**
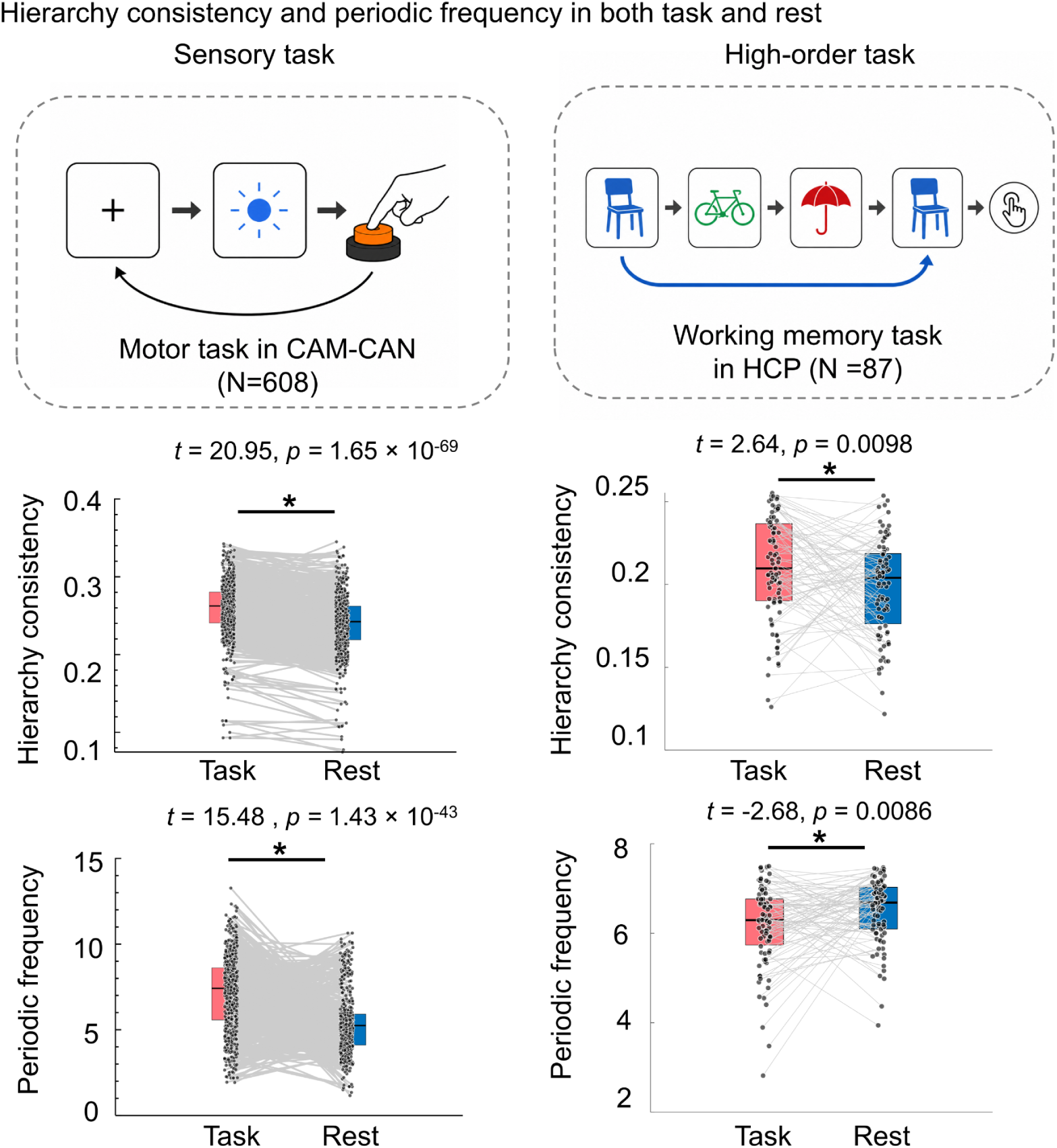
Task-dependent modulation of hierarchy consistency and periodic frequency. **Left,** Cam-CAN sensorimotor task versus rest (*N* = 608). Hierarchy consistency was higher during task than rest (*t* = 20.95, *p* = 1.65 × 10⁻⁶⁹), and periodic frequency was also increased (*t* = 15.48, *p* = 1.43 × 10⁻^43^). **Right,** HCP working-memory task versus rest (*N* = 87). Hierarchy consistency was higher during working memory than rest (*t* = 2.64, *p* = 0.0098), whereas periodic frequency was lower (*t* = −2.68, *p* = 0.0086). Points represent individual participants, grey lines connect paired task and resting-state measurements, and asterisks indicate significant task–rest differences.

Task-derived hierarchy consistency also varied with age. During the task, hierarchy consistency decreased significantly with age (*r* = −0.45, *p* = 1.20 × 10⁻^30^; **Supplementary Fig. 4b**), indicating weaker hierarchy-aligned propagation during task engagement in older adults. Task-derived periodic frequency showed a weaker but significant negative association with age (*r* = −0.15, *p* = 3.23 × 10⁻^5^; **Supplementary Fig. 4b**), suggesting age-related slowing of task-related hierarchy-consistency dynamics. Thus, task engagement consistently strengthened hierarchy-aligned propagation across two independent datasets, whereas the temporal rate of hierarchical reconfiguration was task dependent, increasing during sensorimotor processing but decreasing during working memory.

### Hierarchy consistency and switchboard dynamics vary across human ageing

Having characterized the spectral and spatial properties of hierarchy consistency, we next investigated how these measures varied across the adult lifespan in the Cam-CAN cohort. Hierarchy consistency showed a significant negative association with age (*r* = −0.44, *p* = 2.30 × 10⁻^31^; **Fig. 4a**), indicating that older adults exhibited reduced alignment between moment-to-moment cortical propagation and the macroscale functional hierarchy. The periodic frequency of hierarchy consistency exhibited a significant nonlinear association with age (*r^2^* =0.24, *p* = 1.19 × 10⁻^37^; **Fig. 4a**), suggesting that the rhythmic properties of hierarchy-related propagation dynamics are reorganized across adulthood rather than changing monotonically with age.

**Figure 4.**
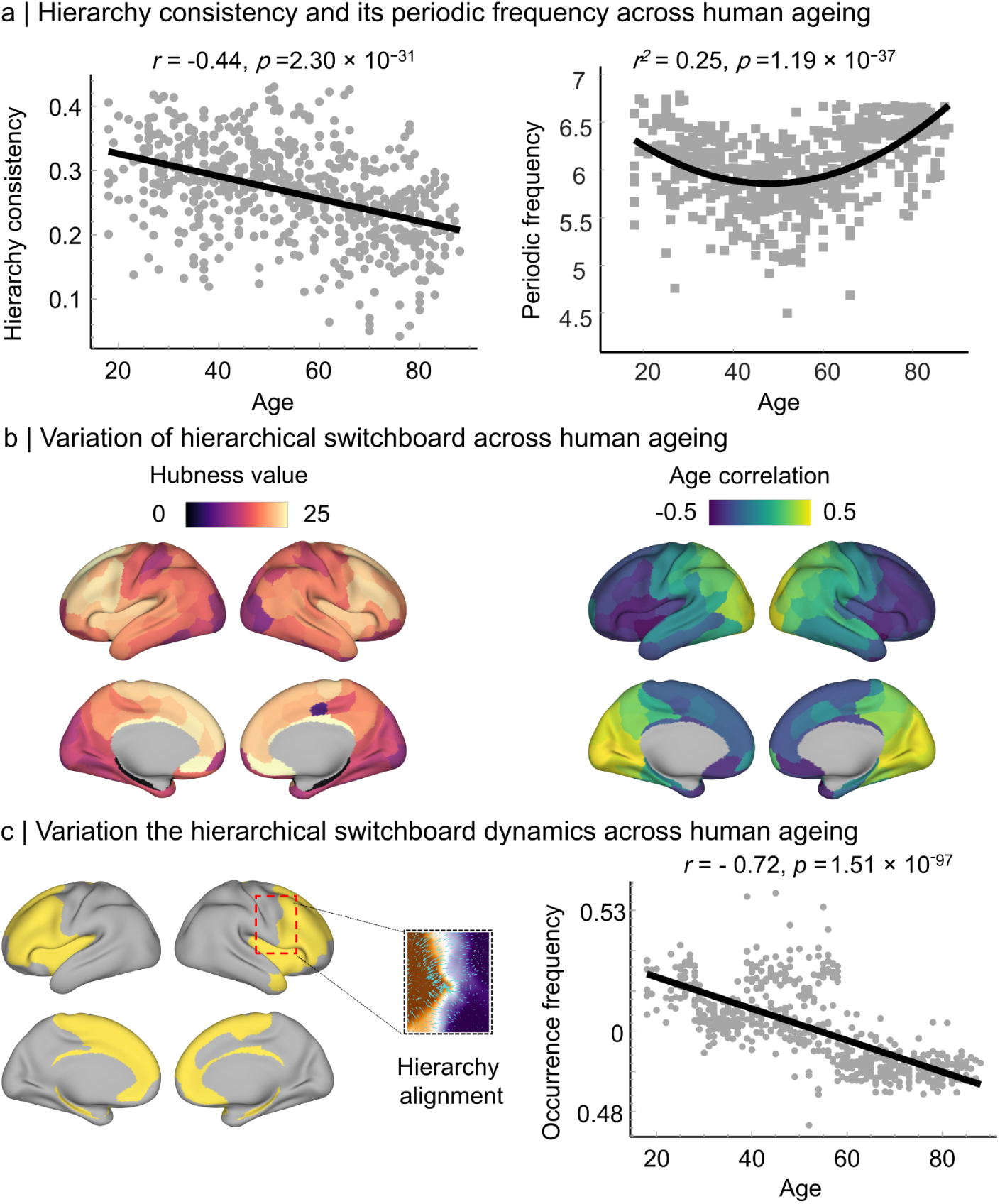
Age-related changes in hierarchy consistency, periodic frequency, and hierarchical switchboard dynamics in CAM-CAN (N=608). **a,** Hierarchy consistency and periodic frequency across adulthood. Hierarchy consistency decreased with age (*r* = −0.44, *p* = 2.30 × 10^-31^). Periodic frequency showed a significant quadratic association with age (*r*² = 0.24, *p* = 1.19 × 10^-37^), with higher values in younger and older adults and lower values in middle age. Each point represents one participant; black lines indicate the fitted models. **b,** Age-related variation in hierarchical switchboard hubness. Left, cortical maps show the group-level distribution of parcel-wise hubness. Right, maps show regional associations between hubness and age, revealing spatially heterogeneous age-related changes across the cortex. **c,** Age-related variation in propagation-state expression within switchboard regions. Yellow masks indicate switchboard-related regions, and the inset illustrates a representative hierarchy-aligned propagation pattern. The occurrence frequency of hierarchy-aligned states decreased with age (*r* = −0.72, *p* = 1.51 × 10⁻⁹⁷). Each point represents one participant, and the black line indicates the linear regression fit.

Age-related effects were also evident in the spatial architecture of the hierarchical switchboard. Group-level hubness maps showed a distributed switchboard organization concentrated in higher-order association cortex, whereas age-correlation maps revealed regionally heterogeneous changes in hubness with ageing (**Fig. 4b**). These patterns indicate that ageing does not uniformly weaken switchboard organization, but instead selectively reshapes the contribution of specific cortical regions to transitions between hierarchy-aligned and hierarchy-opposed propagation states.

We further examined age-related changes in switchboard state dynamics. The occurrence frequency of hierarchy-aligned propagation states within switchboard regions decreased markedly with age (*r* = −0.72, *p* = 1.51 × 10⁻^97^; **Fig. 4c**), indicating that ageing is associated with reduced expression of propagation states aligned with the sensory-to-association hierarchy. Because age-related MEG effects can be influenced by differences in signal quality, source reconstruction, broadband power and data quantity, we further evaluated these associations using sensitivity analyses controlling for available acquisition and signal-quality measures. The age-related effects remained qualitatively consistent after accounting for these covariates and were further supported by robust regression, Spearman rank correlation and leave-one-out analyses. Collectively, these findings suggest that human ageing alters both the strength and temporal organization of hierarchy-consistent propagation.

### Hierarchy consistency and periodic frequency are associated with cognition

Having established that hierarchy-consistency dynamics vary with ageing, we next asked whether individual differences in these measures were behaviourally relevant. Both overall hierarchy consistency and its periodic frequency showed significant positive associations with fluid intelligence (hierarchy consistency: *r* = 0.19, *p* = 1.15 × 10^-6^; periodic frequency: *r* = 0.20, *p* = 7.81 × 10^-7^; **Fig. 5a**) These associations were modest in effect size and should be interpreted with caution given the multifactorial nature of fluid intelligence, but their convergent direction suggests that individuals with stronger and faster hierarchy-related propagation dynamics tended to show higher cognitive performance.

**Figure 5.**
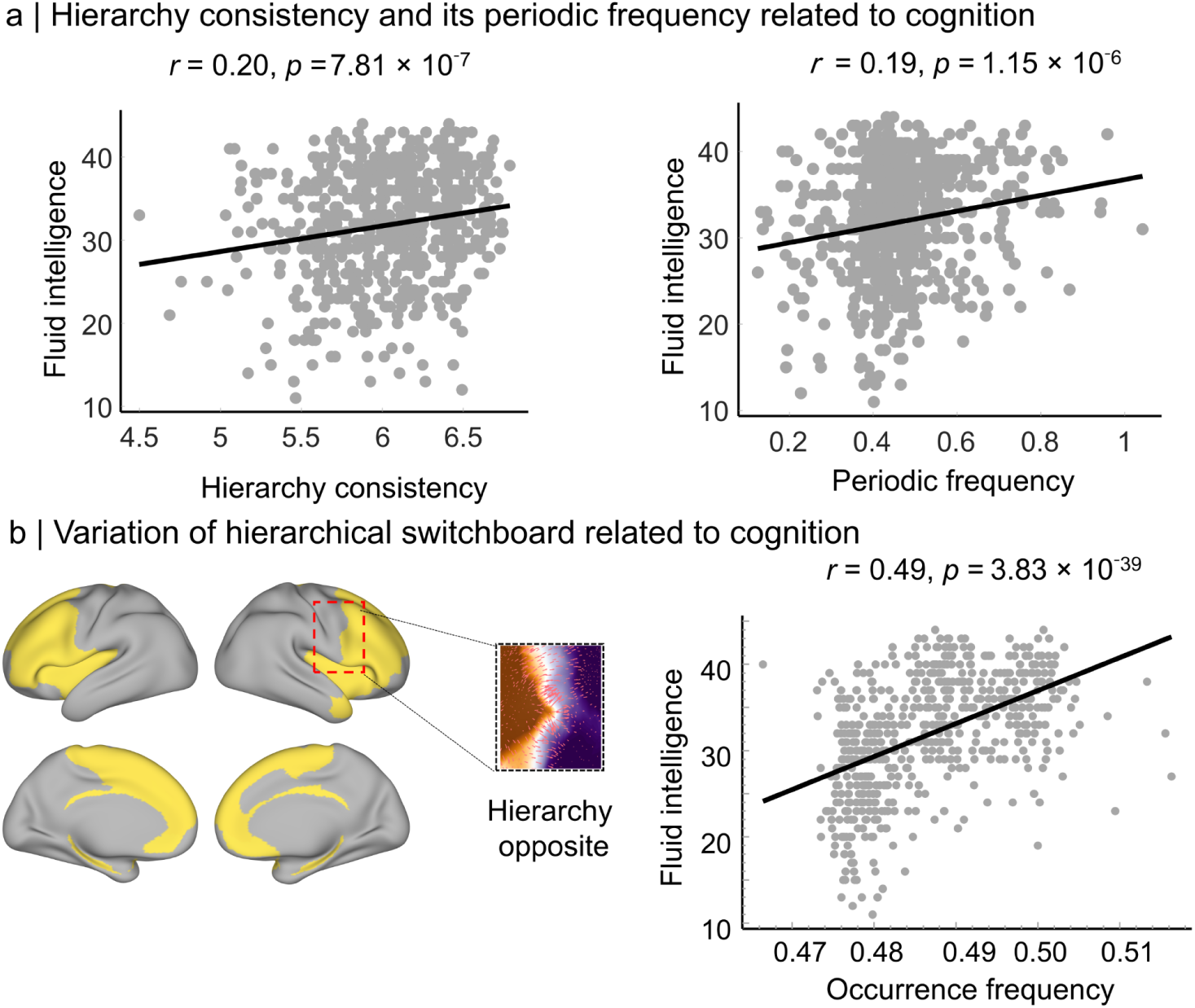
Hierarchy consistency and switchboard dynamics are associated with fluid intelligence in CAM-CAN (N=608). **a,** Associations between hierarchy-consistency dynamics and fluid intelligence. Left, periodic frequency was positively associated with fluid intelligence (*r* = 0.20, *p* = 7.81 × 10⁻⁷). Right, overall hierarchy consistency was also positively associated with fluid intelligence (*r* = 0.19, *p* = 1.15 × 10⁻⁶). Each point represents one participant, and black lines indicate linear regression fits. **b,** Association between hierarchical switchboard dynamics and fluid intelligence. Yellow cortical regions indicate switchboard-related areas identified using *k*-core decomposition, and the inset illustrates a representative hierarchy-opposed propagation state. The occurrence frequency of hierarchy-opposed states within switchboard regions was positively associated with fluid intelligence (*r* = 0.49, *p* = 3.83 × 10⁻^39^). Each point represents one participant, and the black line indicates the linear regression fit.

We next examined whether the cognitive relevance of hierarchy-consistency dynamics extended to the flexibility dimension of switchboard function. Within switchboard-related regions, the occurrence frequency of hierarchy-opposed propagation states was positively associated with fluid intelligence after controlling for age, sex and handedness (*r* = 0.49, *p* = 3.83 × 10⁻^39^; **Fig. 5b**). Given the relatively narrow range of occurrence values, we further assessed robustness using rank-based correlation, robust regression, leave-one-out analysis and permutation testing. The association remained qualitatively consistent across these sensitivity analyses. Together, these results suggest that higher fluid intelligence is associated with both stronger hierarchy-related propagation dynamics and greater relative expression of hierarchy-opposed states within switchboard regions.

### Structural and control architectures constrain the hierarchical switchboard

We next asked whether the spatial organization of the hierarchical switchboard was embedded within established functional and anatomical axes of cortical organization. Regions with different hubness levels occupied systematically distinct positions along the sensory-to-association axis (**Fig. 6a**): lower-hubness regions were concentrated closer to the sensory pole, whereas higher-hubness regions showed broader distributions extending toward the association cortex. This gradient-organized pattern indicates that switchboard participation is not randomly distributed across the cortex, but is structured with respect to macroscale functional architecture.

**Figure 6.**
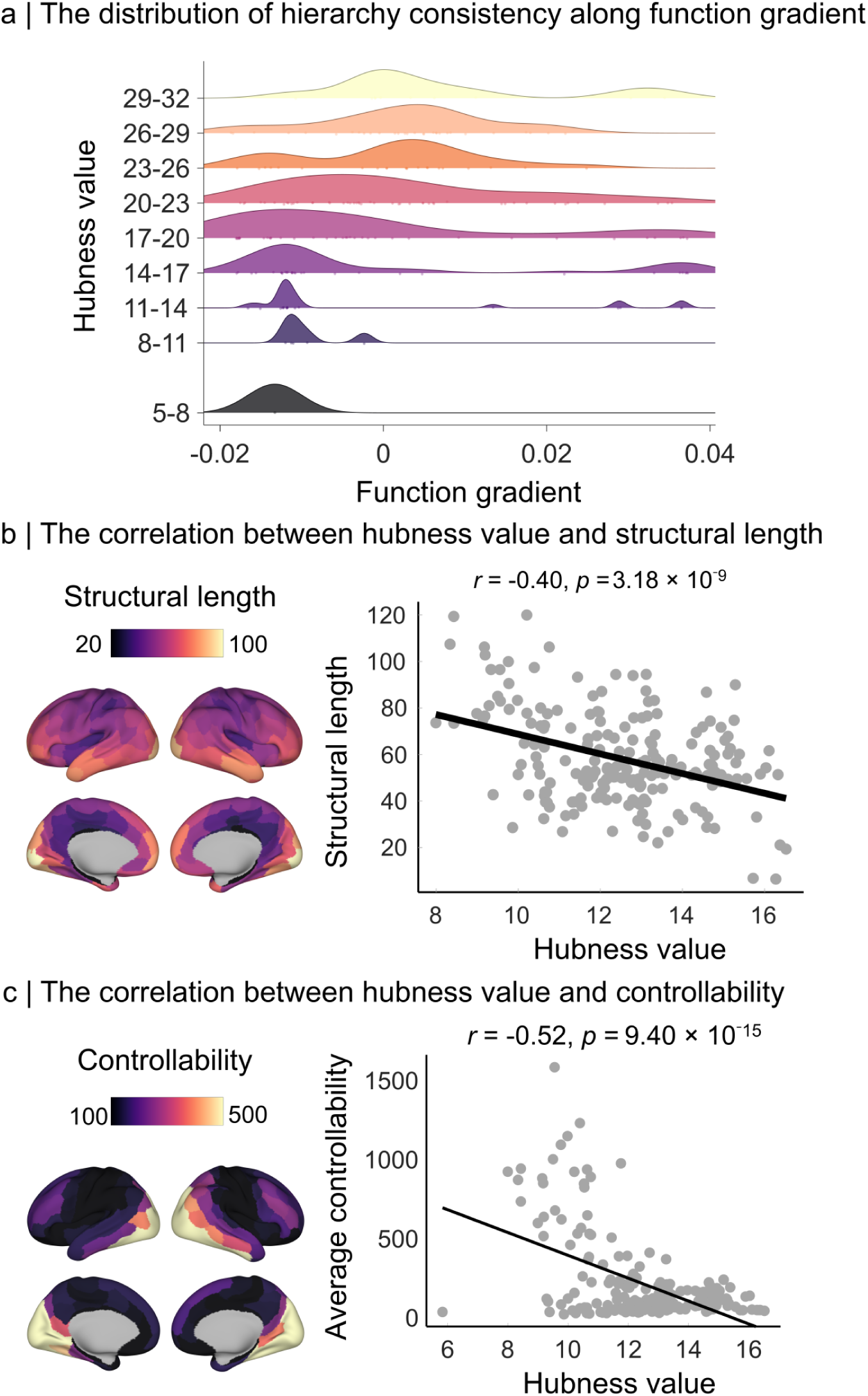
Functional-gradient organization and structural-control constraints of hierarchical switchboard hubness in CAM-CAN (N=608). **a,** Distribution of hierarchical switchboard hubness along the principal cortical functional gradient. Stacked density plots show the distribution of cortical regions across the functional gradient for increasing hubness levels. Higher-hubness regions were shifted toward intermediate-to-high gradient values associated with transmodal association cortex. **b,** Association between switchboard hubness and structural connection length. Left, cortical maps show regional mean structural connection length. Right, hubness was negatively correlated with mean structural connection length (*r* = −0.40, *p* = 3.18 × 10⁻⁹), indicating that regions with higher hubness tended to be associated with shorter structural pathways. Each point represents one cortical parcel, and the black line indicates the linear regression fit. **c,** Association between switchboard hubness and average controllability. Left, cortical maps show regional average controllability derived from the structural connectome. Right, hubness was negatively correlated with average controllability (*r* = −0.52, *p* = 9.40 × 10⁻¹⁵). Each point represents one cortical parcel, and the black line indicates the linear regression fit.

We next asked whether switchboard hubness was related to the structural connectome. Hubness was significantly negatively correlated with mean structural connection length (*r* = −0.40, *p* = 3.18 × 10⁻^9^; **Fig. 6b**), indicating that regions with greater switchboard participation tended to be embedded within shorter anatomical connection pathways. This result suggests that hierarchy-transition hubs may be supported by relatively compact structural wiring. However, because structural length is related to spatial embedding, network membership and connection strength, this association should be interpreted as a structural constraint rather than direct evidence that short pathways causally drive switching.

Finally, we examined whether switchboard hubness was related to average controllability. Hubness was significantly negatively correlated with average controllability (*r* = −0.52, *p* = 9.40 × 10⁻^15^; **Fig. 6c**), indicating that regions with high switchboard participation were less associated with broadly accessible low-energy state transitions. This suggests that switchboard hubs occupy a distinctive position in the network-control landscape, but do not necessarily function as general-purpose control nodes. Complementary analyses further showed that edge-level hierarchy consistency was negatively associated with structural-connectome measures, and regional hierarchy consistency showed a nonlinear association with the principal functional gradient (**Supplementary Fig. 5**). Together, these findings indicate that hierarchical switchboard hubs occupy a functionally and anatomically organized position within the cortical connectome.

### Anatomical frequency relates to hierarchy-consistency rhythm during rest

To examine whether the periodic frequency of hierarchy consistency was related to anatomical constraints, we estimated an anatomical frequency from cortical anatomical decomposition modes. In an example Cam-CAN participant, anatomical decomposition modes were used to derive an anatomical power spectrum, and the dominant anatomical frequency was defined as the frequency corresponding to the largest spectral peak (**Supplementary Fig. 6a**). Across participants, anatomical frequency showed a weak but significant positive association with the periodic frequency of hierarchy consistency during rest (*r* = 0.17, *p* = 9.99 × 10⁻^6^), whereas no significant association was observed during task (*r* = 0.03, *p* = 0.16; **Supplementary Fig. 6b**). These findings suggest that resting hierarchy-consistency rhythms may be partially related to anatomy-derived frequency structure, whereas task engagement may reduce or override this anatomical association. Given the small effect size, this result should be interpreted as exploratory.

## Discussion

In this study, we show that the macroscale sensory-to-association hierarchy provides a dynamic reference axis for human cortical propagation across rest and task states. Using source-resolved MEG combined with Riemannian cortical-flow modelling, we found that hierarchy alignment fluctuated rhythmically during rest, was spatially organized around higher-order association systems, varied across the adult lifespan, and was associated with fluid intelligence. Across behavioural states, hierarchy consistency increased during both sensorimotor and working-memory tasks relative to rest, whereas its periodic frequency was modulated in a task-dependent manner. Together, these findings extend the cortical hierarchy from a predominantly spatial description of cortical organization to a dynamic framework for characterizing the temporal and state-dependent organization of large-scale neural propagation.

A central finding is the identification of a spatially structured hierarchical switchboard that mediates transitions between propagation regimes. Switchboard hubness was preferentially concentrated in higher-order transmodal association systems, including the salience, frontoparietal control, limbic, and default-mode networks — systems well positioned to integrate sensory, cognitive, affective, and executive information streams^2,14,15,19–21^. This spatial organization suggests that these association networks provide the cortical substrate for flexibly switching between bottom-up sensory-to-association propagation and top-down association-to-sensory propagation^3,11,18^. The negative associations between switchboard hubness, structural connection length, and average controllability further demonstrate that hierarchy switching is not an arbitrary or unconstrained process, but is tightly regulated by anatomical wiring economy and network-control architecture^22–24^. Switchboard regions thus appear to coordinate flexible propagation-regime transitions through compact structural circuits while occupying a distinctive and constrained position in the cortical control landscape — one that prioritizes selective state-switching over broad, energetically cheap state-transition capacity.

These findings extend and recontextualize prior work on cortical gradients and neural propagation in several important respects^1,18,25–27^. Previous studies have firmly established the sensory-to-association hierarchy as a major spatial axis of cortical organization, with expressions spanning functional connectivity, cortical microstructure, myeloarchitecture, receptor density profiles, and patterns of cognitive specialization^3,4,25,28^. Our framework adds a critical dynamic dimension to this spatial picture: by combining MEG source imaging, cortical-flow estimation, and gradient-based directional alignment, we demonstrate that hierarchy-related propagation is a time-varying process with identifiable rhythmic periodicity and switchboard-like spatial organization. This provides a principled conceptual and methodological bridge between static spatial models of cortical hierarchy and temporal models of large-scale neural dynamics, opening new avenues for characterizing the temporal dimension of cortical gradient organization.

Hierarchy-consistency dynamics were further shown to be sensitive to both intrinsic ageing processes and momentary cognitive state, demonstrating their function across the adult lifespan. Across ageing, hierarchy consistency declined progressively, and hierarchy-aligned propagation states became significantly less frequent within switchboard regions — a pattern indicative of an age-related shift in the balance between propagation regimes, rather than a global or uniform reduction in cortical dynamics^4,11,29^. This selective reorganization may reflect differential age-related vulnerability of transmodal association systems, which are known to exhibit pronounced structural and functional decline with advancing age^30^. Task engagement, by contrast, consistently increased hierarchy consistency in both the Cam-CAN sensorimotor task and the HCP working-memory task, indicating stronger alignment of cortical propagation with the sensory-to-association hierarchy. However, periodic frequency was modulated in a task-specific manner, increasing during sensorimotor processing but decreasing during working memory. These findings suggest that task engagement strengthens hierarchy-consistent propagation while flexibly adjusting its temporal scale to behavioural demands. The persistence of negative age associations during task further suggests that ageing may specifically impair the brain’s capacity to sustain and recruit hierarchy-aligned flow under cognitive demands — a deficit that may contribute to age-related declines in cognitive efficiency and flexibility^31,32^.

The behavioural associations observed in this study indicate that hierarchy-consistency dynamics carry meaningful cognitive information at the individual level. Individuals with stronger hierarchy consistency and faster periodic frequency showed higher fluid intelligence, demonstrating that the fidelity and tempo of hierarchy-related propagation dynamics reflect aspects of cognitive capacity that generalize beyond age effects^31^. Critically, greater occurrence of hierarchy-opposed propagation states within switchboard regions was associated with substantially higher fluid intelligence — an effect size notably larger than those observed for overall hierarchy consistency or periodic frequency alone. This dual association suggests that efficient cognition does not simply reflect maximized alignment with the cortical hierarchy; rather, it requires an optimized dynamic balance between stable, hierarchy-consistent information routing and flexible, context-sensitive departures from it^14,33^. The capacity to transiently enter hierarchy-opposed propagation states — potentially reflecting top-down modulation, predictive processing, or internally guided cognition — may therefore represent a critical component of the neural architecture underlying fluid cognitive capacity^14^.

Several limitations should be considered. First, cortical-flow estimates derived from source-resolved MEG reflect apparent macroscopic propagation of electromagnetic field activity and cannot directly establish synaptic-level communication or causal transmission between cortical regions. Second, hierarchy consistency quantifies angular alignment between cortical-flow vectors and a functional-gradient reference axis; it should therefore be interpreted as a measure of propagation alignment rather than direct information flow. Finally, behavioural associations with fluid intelligence were correlational, and several effects were modest in size; these results should therefore be interpreted as evidence of behavioural relevance rather than direct cognitive causality.

## Conclusion

Human cortical propagation is dynamically organized by the sensory-to-association functional hierarchy. By quantifying hierarchy consistency from source-resolved MEG, we identify a rhythmic, switchboard-supported organization that coordinates transitions between hierarchy-aligned and hierarchy-opposed propagation states. This dynamic architecture is modulated by task demands, reorganized across ageing, and associated with individual differences in fluid intelligence. Together, these findings establish hierarchy consistency as a framework for linking macroscale cortical organization with the temporal dynamics, flexibility, and behavioural relevance of large-scale neural propagation.

## Materials and Methods

### Participants and datasets

Primary analyses were performed using resting-state data from the Cambridge Centre for Ageing and Neuroscience (Cam-CAN) cohort^34^. The final sample included 608 healthy adults aged 18–88 years (307 male, 301 female; mean age, 54.19 ± 18.19 years), all of whom had resting-state MEG and T1-weighted MRI data. MEG was acquired during eyes-closed rest using a whole-head MEGIN system (Helsinki, Finland) at a sampling rate of 1,000 Hz. Fluid intelligence was quantified using the Cam-CAN composite score (mean ± s.d., 31.80 ± 6.79), derived from four subtests: series completion, classification, matrices and conditions. Resting-state functional MRI data from the Cam-CAN cohort were used to derive the cortical functional hierarchy against which MEG propagation was referenced. Resting-state fMRI was acquired using gradient-echo echo-planar imaging with a repetition time of 1,970 ms, echo time of 30 ms, flip angle of 78°, voxel size of 3 × 3 × 4.44 mm³, and 32 axial slices. Each resting-state fMRI run lasted approximately 8 min 40 s.

Replication analyses were conducted using resting-state data from the Open MEG Archive (OMEGA) dataset^35^. After quality control and assessment of data completeness, the replication sample comprised 83 healthy adults (40 female, 43 male; mean age, 28.66 ± 6.97 years). MEG data were acquired using a 275-channel whole-head CTF system (Port Coquitlam, British Columbia, Canada) at a sampling rate of 2,400 Hz. Unless otherwise specified, the same preprocessing pipeline, analysis procedures and metric definitions were applied to both datasets. Both repositories contain de-identified data collected under the ethical approvals and consent procedures of the original studies.

We analysed resting-state and working-memory MEG data from 87 healthy adults (40 females, 47 males; age 18–40 years) in the WU–Minn HCP Young Adult dataset. Participants without usable MEG or corresponding structural MRI data were excluded. MEG was acquired using a 248-channel MAGNES 3600 system at a sampling rate of 2035 Hz. Resting-state recordings comprised ∼6-min eyes-open runs, and task recordings included a visual N-back working-memory paradigm with alternating 0-back and 2-back conditions (see Supplementary Material for details).

### MEG preprocessing and source reconstruction

Preprocessing was performed in Brainstorm^36^ in accordance with established recommendations for MEG reporting ^37^. Continuous data were band-pass filtered between 0.3 and 200 Hz using finite impulse response filters. Line noise and its harmonics were attenuated using notch filters at 50, 100 and 150 Hz for Cam-CAN, and at 60, 120 and 180 Hz for OMEGA and HCP. Bad channels and noisy data segments were identified and excluded. Ocular and cardiac artifacts were reduced using signal-space projection, with projectors estimated from EOG- and ECG-locked epochs, respectively.

For each participant, power spectral density was estimated using Welch’s method with 2-s windows and 50% overlap. To define an adaptive broadband upper frequency limit, we identified the frequency below which 95% of the total spectral power was contained. Clean data was then divided into non-overlapping 30-s epochs, a window length that provides stable estimates of resting-state activity. Each epoch was resampled at four times the participant-specific cutoff frequency to minimize aliasing and reduce computational demands.

Structural MRI data were processed with FreeSurfer to reconstruct individual cortical surfaces and triangular meshes^38^. MEG and MRI data were co-registered using digitized scalp points. Forward models were computed using overlapping-spheres head models, and source time series were reconstructed with linearly constrained minimum-variance beamforming with depth-bias correction. Noise covariance matrices were estimated from empty-room recordings. Source-resolved time series were analysed for broadband activity and for canonical frequency bands.

### Cortical-flow estimation on a Riemannian surface

We estimated cortical propagation by modelling source-reconstructed MEG activity as a time-varying scalar field on the cortical surface. The cortex was represented as a triangular mesh approximating a two-dimensional Riemannian manifold embedded in three-dimensional space. Local cortical-flow vectors were estimated using an optical-flow formulation adapted to the cortical manifold, in which temporal changes in activity were related to spatial gradients on the surface under a local conservation assumption^39^. To address ambiguity in local flow estimation, we minimized an energy functional that combined fidelity to the observed temporal dynamics with spatial smoothness of the vector field. Individual cortical surfaces and flow fields were registered to the fsaverage spherical template to ensure vertex-wise correspondence across participants. Flow vectors were then projected onto an anatomy-informed local tangent-plane reference frame, with directions expressed relative to posterior–anterior and superior–inferior axes, enabling quantification of cortical propagation direction in a common anatomical coordinate system^11^.

### fMRI preprocessing and functional gradient construction

Raw fMRI data were converted to BIDS format^40^ using HeuDiConv and preprocessed with fMRIPrep^41^. Preprocessing included standard anatomical reconstruction, spatial normalization, slice-timing and motion correction, co-registration to each participant’s T1-weighted image, and minimal nuisance regression to reduce non-neuronal confounds while preserving individual functional architecture. To enable direct comparison with MEG-derived cortical dynamics, preprocessed fMRI data were resampled to the same 15,002 cortical locations used for the MEG source representation. Functional connectivity matrices were computed using zero-lag Pearson correlations between cortical time series. Individual functional gradients were then estimated with BrainSpace^42^ by applying Fisher z-transformation, proportional thresholding of the strongest 10% of connections, cosine-kernel affinity estimation and diffusion map embedding^43^. The first three gradients were retained and aligned to a group-level reference using Procrustes alignment.

### Definition of hierarchy consistency

To quantify the alignment between cortical propagation and the macroscale functional hierarchy, we compared the local cortical-flow direction with the local direction of the principal functional gradient^18^. At each cortical vertex, the local direction of the sensory-to-association hierarchy was defined as the surface gradient of the principal functional-gradient map and normalized to yield a unit vector field. For each vertex and time point, we calculated the unsigned angular difference (Δtheta) between the cortical-flow vector and the local hierarchy direction, bounded between (0°) and (180°).

Flow was classified as hierarchy-aligned propagation when (0° ≤ Δθ ≤ 90°), indicating propagation along the sensory-to-association direction. Flow was classified as hierarchy-opposed propagation when (90° < Δθ ≤ 180°), indicating propagation opposite to the functional hierarchy. For each time point, hierarchy consistency was defined as the proportion of cortical vertices exhibiting hierarchy-aligned propagation, that is, vertices for which the angular difference between the cortical-flow vector and the local functional-gradient direction satisfied 0° ≤ Δθ ≤ 90°. Specifically, hierarchy consistency(t) = N_aligned(t) / N_total, where N_aligned(t) denotes the number of cortical vertices classified as hierarchy-aligned at time t, and N_total denotes the total number of cortical vertices^11^. Thus, hierarchy consistency ranges from 0 to 1, with higher values indicating that a larger proportion of the cortex exhibits propagation aligned with the sensory-to-association hierarchy.

### Extraction of the periodic frequency of hierarchy consistency

To quantify the temporal periodicity of hierarchy-consistent propagation, we estimated the spectral structure of the hierarchy-consistency time series. For each participant, the power spectral density of the hierarchy-consistency signal was first computed and spectrally parameterized to separate periodic components from the aperiodic (1/f)-like background. Spectral parameterization was performed in log–log power–frequency space using a fixed aperiodic model^44^. The estimated aperiodic background was then subtracted from the original power spectrum to obtain an aperiodic-adjusted spectrum, enabling the identification of periodic peaks independently of broadband spectral decay. The dominant periodic component was defined as the peak with the largest aperiodic-adjusted power within the analysed frequency range (2-30 Hz). The centre frequency of this peak was extracted as the individual periodic frequency of hierarchy consistency. This frequency indexes the characteristic timescale of rhythmic fluctuations in the prevalence of hierarchy-aligned cortical propagation.

To ensure that the extracted periodic frequency of hierarchy consistency did not simply reflect the aperiodic 1/f-like background, we evaluated the goodness-of-fit of the spectral parameterization model for each participant. Fitting scores were summarized in both the Cam-CAN and OMEGA datasets. We also inspected individual power spectra and group-average spectra to confirm that the identified periodic component was extracted from the aperiodic-adjusted spectrum rather than from the raw low-frequency power alone.

### Identification of hierarchical switchboard hubs

To identify cortical regions that coordinated transitions between hierarchy-aligned and hierarchy-opposed propagation states, we constructed an inter-regional synchronization network from parcel-wise hierarchy-related temporal profiles. For each participant, hierarchy-related signals were extracted for each of the 200 Schaefer cortical parcels. Pairwise phase-locking values (PLVs) were calculated between parcel-wise hierarchy-related temporal profiles, yielding a 200 × 200 synchronization matrix for each participant.

To identify regions embedded within the core of this hierarchy-transition network, individual connectivity matrices were thresholded using a proportional threshold, retaining the strongest 10% of connections. The resulting sparse matrices were then entered into k-core decomposition. Briefly, each weighted matrix was converted into a binary undirected adjacency matrix and symmetrized. Nodes with degree lower than the current k level were iteratively removed, and each parcel was assigned a coreness value corresponding to the highest k-core in which it remained. Higher coreness therefore indicated that a cortical parcel was more centrally embedded within the synchronized hierarchy-transition network, suggesting stronger involvement in coordinating transitions between hierarchy-aligned and hierarchy-opposed propagation states.

Group-level switchboard hubness maps were generated by averaging parcel-wise coreness values across participants. To characterize the network-level organization of hierarchical switchboard hubs, coreness values were summarized within canonical functional systems, including visual, somatomotor, dorsal attention, salience/ventral attention, limbic, frontoparietal/control and default-mode networks. Regions with high coreness were interpreted as hierarchical switchboard hubs supporting coordinated alternation between propagation aligned with and opposed to the macroscale functional hierarchy.

### Switchboard state dynamics

To characterize the temporal dynamics expressed within hierarchical switchboard hubs, we quantified the occurrence of hierarchy-aligned and hierarchy-opposed propagation states within regions identified by the group-level switchboard hubness map. Switchboard-related regions were defined as parcels with high coreness in the hierarchy-transition network, reflecting their central embedding within synchronized hierarchy-angle dynamics. For each participant, vertex- or parcel-wise propagation states were first classified according to their angular alignment with the principal functional hierarchy. We then calculated, within switchboard-related regions, the proportion of time points assigned to hierarchy-aligned and hierarchy-opposed propagation states. These occurrence-frequency measures were used to assess whether hierarchical switchboard hubs preferentially expressed propagation along the sensory-to-association hierarchy, propagation in the opposite direction, or a balanced temporal alternation between the two regimes. In this way, switchboard state dynamics captured the extent to which core regions of the hierarchy-transition network supported flexible switching between aligned and opposed modes of cortical propagation.

### Structural length and network controllability

Diffusion-weighted imaging data from Cam-CAN were preprocessed with QSIPrep^45^, and weighted structural connectomes were reconstructed using a multi-shell, multi-tissue anatomically constrained tractography workflow with SIFT2 streamline reweighting^46^. To assess whether hierarchical switchboard regions were constrained by structural wiring, we related regional switchboard hubness to two connectome-derived measures: anatomical connection length and average controllability. Regional structural length was defined as the mean length of structural connections incident on each cortical region. Average controllability was computed within a linear network control framework, using the normalized structural connectivity matrix to estimate the ability of each region to drive the brain toward nearby, low-energy states^24,47^. Regional hubness values were then correlated with structural length and average controllability to determine whether hierarchy-switching regions occupy distinctive positions in the brain’s anatomical and control-energy architecture.

### Age effect

Age associations were tested using correlation and regression analyses. For resting-state data, we examined associations between age and hierarchy consistency, periodic frequency, switchboard hubness and switchboard state occurrence, with sex and handedness included as covariates. To evaluate whether age-related associations were robust to potential data-quality confounds, we repeated the main regression models with additional covariates where available, including head-motion estimates, the number of retained clean epochs, global source power and other signal-quality indicators. Robust regression, Spearman rank correlation and leave-one-out analyses were further used to assess the sensitivity of age effects to outliers.

For task data, hierarchy consistency and periodic frequency were computed using the same procedures as in resting-state data, and paired comparisons were used to test differences between task and resting-state conditions. Age associations in task-derived hierarchy consistency and periodic frequency were then evaluated using correlation and regression analyses, with sex and handedness included as covariates. Where available, the same data-quality covariates and outlier-sensitivity analyses were applied to task-derived measures. Statistical significance was assessed using permutation testing, and multiple comparisons were controlled using false-discovery-rate correction where appropriate.

### Cognitive relevance

To assess behavioural relevance, fluid intelligence scores were related to hierarchy consistency, periodic frequency and switchboard state occurrence, with age, sex and handedness included as covariates. These analyses tested whether individuals with stronger hierarchy-aligned propagation, faster hierarchy-consistency rhythms or greater flexibility in hierarchy-opposed switchboard states showed better cognitive performance. To determine whether cognitive associations were independent of lifespan effects, age was included as a covariate in all cognitive models. Where available, we additionally controlled for data-quality measures, including head-motion estimates, the number of retained clean epochs, global source power and signal-quality indicators. Robust regression, Spearman rank correlation and leave-one-out analyses were used to evaluate whether cognitive associations were driven by outliers. Statistical significance was assessed using permutation testing, and multiple comparisons were controlled using false-discovery-rate correction where appropriate.

### Hierarchy consistency in task

Task-state MEG data were obtained from the Cam-CAN sensorimotor task, in which participants detected brief auditory, visual or audiovisual stimuli and responded with a right index-finger button press. MEG recordings were epoched relative to the button press to capture movement-related cortical dynamics. After trial- and participant-level quality control, source-level task epochs were used to quantify broadband cortical flow. Hierarchy consistency and its periodic frequency were computed using the same procedures as in resting-state data, allowing us to test whether hierarchy-dependent propagation observed at rest was preserved or reorganized during externally cued sensorimotor behaviour. Task–rest differences were assessed with paired comparisons, and age associations in task-derived measures were evaluated using correlation analyses. In the HCP dataset, task-state MEG data were obtained from the visual N-back working-memory paradigm comprising alternating 0-back and 2-back conditions. Source-resolved working-memory and resting-state MEG data were analysed using the same cortical-flow framework, and hierarchy consistency and its periodic frequency were calculated using the same definitions as in the primary analyses. These complementary task datasets allowed us to test whether hierarchy-dependent propagation was flexibly reorganized across externally driven sensorimotor and higher-order working-memory demands. Task–rest differences were assessed using paired comparisons.

## Data availability

The resting-state MEG, structural MRI, diffusion MRI and resting-state fMRI data used for the primary analyses are available through the Cambridge Centre for Ageing and Neuroscience (Cam-CAN: https://www.cam-can.org/). Resting-state MEG data used for independent replication are available through the Open MEG Archive (OMEGA: https://www.mcgill.ca/bic/neuroinformatics/omega). Resting-state fMRI data from the Human Connectome Project (HCP: https://www.humanconnectome.org/) were used to construct the group-level functional-gradient reference template. Resting-state and working-memory MEG data from the HCP Young Adult dataset were additionally used for independent task validation. Cortical functional gradients were computed using the BrainSpace toolbox (https://brainspace.readthedocs.io/en/latest/).

## Code availability

Analysis code supporting the findings of this study is available at https://github.com/Laoma29/Cortical-Flow-Project.

## Acknowledgements

X.L. was supported by the China Scholarship Council.

## Supplementary Materials

### Cortical-flow estimation on a Riemannian surface

We modelled the cortical surface as a two-dimensional Riemannian manifold embedded in three-dimensional space and represented it computationally as a triangular mesh. Cortical activity was treated as a time-varying scalar field defined on this manifold.

In local coordinates, the surface normal at each vertex was defined by the cross-product of the partial derivatives of the embedding function:

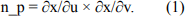

Because this normal is determined solely by the local geometry of the surface, it is invariant to the specific coordinate parametrization. Let I denote the scalar field representing cortical activity, such as the source-reconstructed MEG signal. Its differential maps tangent vectors on the cortical manifold to real values and quantifies the local directional variation required to define spatial gradients and cortical flow:

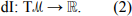

For tangent vectors v1 and v2, the local differential interaction can be written as

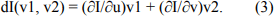

This formulation provides the geometric quantities required to estimate local propagation directly on the cortical surface.

Cortical flow is conceptually analogous to optical flow in computer vision, in which apparent motion is inferred from successive image frames (Lefèvre and Baillet, 2008). Here, cortical activity was considered as a sequence of scalar maps evolving on the cortical manifold. Assuming local conservation of this scalar field along flow trajectories, the cortical-flow field satisfies the transport equation

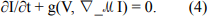

Here, g(·,·) denotes the Riemannian metric, which defines the local inner product on the curved cortical surface, and ∇ℳI is the surface gradient of the scalar field (Lefèvre and Baillet, 2008). This conservation assumption was used as a local approximation at MEG timescales, over which activity patterns vary smoothly across consecutive samples.

As in classical optical-flow estimation, the transport equation alone does not uniquely determine the flow field in regions with locally ambiguous gradients, reflecting the aperture problem. We therefore estimated the cortical-flow vector field by minimizing an energy functional that balances fidelity to the observed temporal dynamics with spatial smoothness:

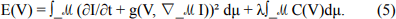

In this expression, dμ is the Riemannian volume form, which determines the local integration weights on the cortical manifold; C(V) penalizes spatially abrupt variation in the vector field; and λ controls the trade-off between data fidelity and smoothness. We set λ = 0.01^6^.

### Anatomy-informed geodesic reference frame

Individual cortical surfaces and their associated flow fields were registered to the fsaverage spherical template to establish vertex-wise correspondence across participants. To quantify propagation direction in an anatomically interpretable coordinate system, we constructed a local tangent-plane reference frame at each cortical vertex. Within each hemisphere, the posterior–anterior axis was defined in template space using the geodesic direction connecting the most posterior and anterior cortical vertices. A superior–inferior axis was then obtained as the orthogonal direction within the local tangent plane, yielding a right-handed anatomical coordinate frame at each vertex. Hemisphere-specific sign conventions were applied to ensure that vector orientations were interpreted consistently across medial and lateral cortical surfaces. Flow vectors were projected onto this local anatomical frame and converted to angular directions. Under this convention, 90° corresponded to posterior-to-anterior propagation, whereas 270° corresponded to anterior-to-posterior propagation.

### fMRI preprocessing and functional gradient construction

Raw fMRI data were converted from DICOM to BIDS format using HeuDiConv v0.13.1 and preprocessed with fMRIPrep v23.0.2. Structural images underwent bias-field correction, skull stripping, tissue segmentation, cortical surface reconstruction and spatial normalization to MNI space. Functional images were corrected for slice timing and head motion and co-registered to each participant’s T1-weighted anatomical image. A minimal denoising procedure was applied to reduce non-neuronal confounds while preserving individual functional architecture, including high-pass filtering, regression of motion parameters and tissue-derived nuisance signals, linear detrending and z-score normalization.

To facilitate direct comparison with the MEG source space, preprocessed fMRI data were resampled to 15,002 cortical locations, matching the dimensionality of the MEG source representation. Individual functional connectivity matrices were then constructed by computing zero-lag Pearson correlations between regional time series. Functional gradients were estimated using BrainSpace. For each participant, connectivity matrices were Fisher z-transformed and thresholded by retaining the strongest 10% of connections for each cortical location. A cosine-similarity kernel was then used to construct an affinity matrix, and diffusion map embedding was applied to derive low-dimensional representations of functional connectivity organization. The first three gradients were retained for subsequent analyses and aligned to a group-level reference template using Procrustes alignment.

### Structural length and network controllability

Diffusion-weighted imaging data from Cam-CAN were preprocessed with QSIPrep v0.17.0. Preprocessing included denoising, Gibbs-ringing correction, bias-field correction, intensity normalization, correction for head motion and eddy-current distortions, and registration to standard space. Structural connectomes were reconstructed using the mrtrix_multishell_msmt_ACT-hsvs workflow. Fibre-orientation distributions were estimated using multi-shell multi-tissue constrained spherical deconvolution, followed by probabilistic tractography with the iFOD2 algorithm and anatomically constrained tractography informed by T1-weighted tissue segmentations. Streamlines were subsequently reweighted with SIFT2, yielding weighted structural connectivity matrices for downstream analyses.

To test whether hierarchical switchboard regions were constrained by anatomical wiring and network-control architecture, we related regional switchboard hubness to two structural-connectome-derived measures: connection length and average controllability. For each participant, the weighted structural connectivity matrix was used to estimate inter-regional anatomical connection length. Regional structural length was summarized as the mean length of all structural connections incident on a given cortical region. We then correlated regional hubness values with structural length to determine whether hierarchy-switching hubs were preferentially embedded within shorter or longer anatomical pathways.

Average controllability was computed following the network control theory framework introduced by Gu et al. In this framework, the brain is modelled as a discrete-time linear dynamical system in which the pattern of future brain activity is constrained by the underlying structural connectome:

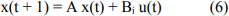

where x(t) denotes the brain state at time t, defined as the vector of regional neural activity across N brain regions; A is the weighted structural connectivity matrix, normalized to ensure stable dynamics; Bᵢ is an input matrix specifying region i as the control node; and u(t) represents the external input applied to that region. For each cortical region, we estimated its ability to control the system by computing the controllability Gramian,

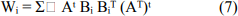

over an infinite time horizon for the stable system. Average controllability was defined as trace(Wᵢ), which quantifies the extent to which input to region i can move the brain into many nearby, easily reachable states with relatively low energy. Regions with high average controllability are therefore interpreted as structural-control hubs that can broadly influence low-energy transitions across the brain’s dynamical state space.

Regional average controllability values were correlated with switchboard hubness to test whether cortical regions that participate strongly in hierarchy switching occupy distinctive positions in the control-energy landscape. A positive association would indicate that switchboard hubs are preferentially located in regions capable of driving the system toward many easily reachable states, whereas a negative or null association would suggest that hierarchy switching is not primarily explained by average low-energy controllability.

### Anatomical frequency estimation

To test whether the periodic frequency of hierarchy-consistency dynamics was related to intrinsic anatomical constraints, we estimated an anatomy-derived frequency for each participant. This analysis was motivated by recent eigenmode-based neural field models, which suggest that large-scale brain activity can be decomposed into spatial eigenmodes of the cortical surface and that different eigenmodes are associated with distinct characteristic frequencies. In this framework, cortical geometry provides a modal basis for brain activity, and the eigenvalues of these modes constrain the frequency structure of large-scale neural rhythms.

For each participant, the cortical surface was represented as a triangular mesh, and anatomical decomposition modes were obtained by solving the Laplace–Beltrami eigenvalue problem on the cortical manifold. These modes provide an ordered set of spatial basis functions determined by cortical geometry, with lower-order modes representing broad spatial patterns and higher-order modes representing finer spatial variation. Participant-specific cortical activity or hierarchy-consistency-related spatial patterns were projected onto these anatomical modes to obtain a modal power spectrum. The dominant anatomical frequency was defined as the frequency corresponding to the largest peak in this anatomical power spectrum.

We then tested whether this anatomy-derived frequency was associated with the empirically observed periodic frequency of hierarchy consistency during rest and task. Correlations were computed across participants separately for resting-state and task conditions.

### Hierarchy consistency in motor task

Task-state MEG data were obtained from the Cambridge Centre for Ageing and Neuroscience dataset. We used the sensorimotor task recordings, in which participants performed a simple cued button-press paradigm. During the task, participants were presented with brief auditory, visual, or audiovisual stimuli and were instructed to respond with a right index-finger button press whenever a stimulus was detected. The task consisted predominantly of bimodal audiovisual trials, together with a small number of unimodal auditory or visual trials included to discourage strategic attention to a single sensory modality. Stimulus onsets were pseudorandomly ordered with variable stimulus-onset asynchronies, thereby providing temporally separated task events suitable for estimating movement-related neural dynamics.

Task MEG recordings were segmented into event-related epochs time-locked to the button press. This procedure allowed us to isolate task-evoked sensorimotor dynamics surrounding movement execution. Trials with delayed responses or insufficient separation from the preceding response were excluded to reduce contamination from poor task performance or overlapping event-related activity. For each retained participant, cleaned task epochs were used to estimate task-state cortical dynamics. Source-level time series were extracted from cortical regions and used to characterize task-evoked broadband activity. In line with prior work on Cam-CAN sensorimotor MEG data, broadband dynamics were treated as transient burst-like events rather than sustained oscillatory activity. We therefore focused on the timing and spatial organization of broadband activity around movement, allowing us to quantify how task engagement reorganizes large-scale cortical activity relative to resting state.

This task-state dataset was used to test whether cortical hierarchy-related dynamics observed during rest were preserved, amplified, or reorganized during externally cued sensorimotor behavior. By comparing task-state dynamics with resting-state activity from the same dataset, we assessed whether hierarchy-dependent cortical flow reflects a general intrinsic organizational principle or becomes selectively structured under task demands.

### Human Connectome Project MEG dataset

Magnetoencephalography (MEG) and structural MRI data were obtained from the WU–Minn Human Connectome Project (HCP) Young Adult dataset. The present analysis included 87 healthy adults with usable MEG recordings and corresponding anatomical MRI data. MEG recordings were acquired using a whole-head MAGNES 3600 system (4D Neuroimaging, San Diego, CA, USA) equipped with 248 magnetometer channels and 23 reference channels. Continuous MEG signals were sampled at 2035 Hz, with simultaneously recorded electrooculography and electrocardiography channels for monitoring ocular and cardiac activity. Participants underwent both eyes-open resting-state MEG and task MEG sessions, including a visual N-back working-memory paradigm.

Resting-state MEG was acquired over approximately 6 min each. Participants were instructed to remain still and relaxed with their eyes open while fixating on a centrally presented red crosshair against a dark background. Electrocardiography and electrooculography were recorded simultaneously for physiological artifact monitoring. Resting-state recordings preceded the task of MEG acquisitions.

Working-memory processing was assessed using a visual N-back paradigm comprising alternating 0-back and 2-back conditions. During 2-back blocks, participants indicated whether the current stimulus matched the stimulus presented two trials previously, thereby requiring continuous updating and maintenance of information in working memory. During 0-back blocks, participants responded when the current stimulus matched a predefined target. Each task block consisted of 10 trials lasting 2.5 s each, including a 2-s stimulus presentation followed by a 500-ms inter-trial interval. Task blocks were interleaved with fixation blocks.

---

**Supplementary Fig. 1.**
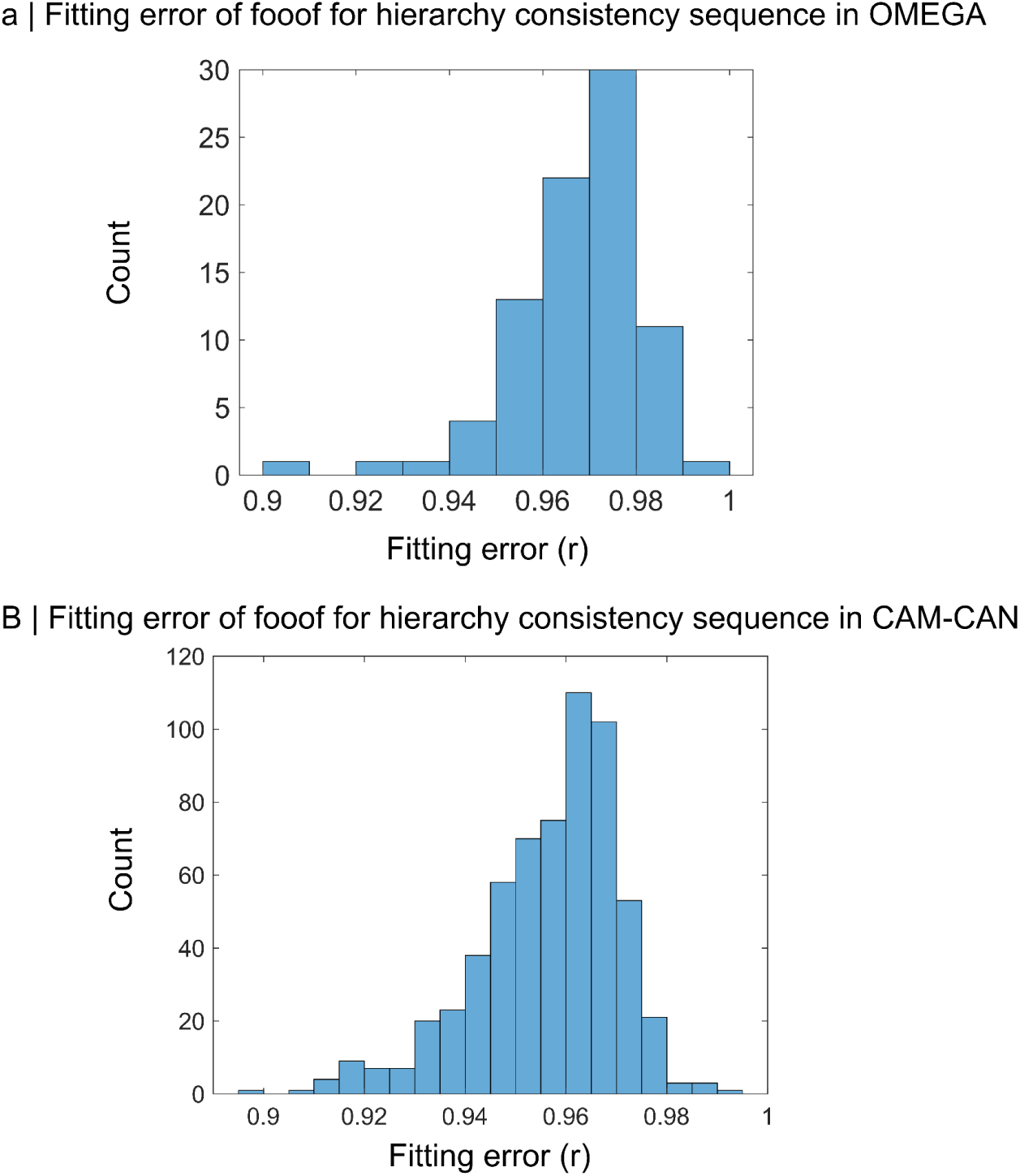
Goodness-of-fit for spectral parameterization of hierarchy-consistency dynamics. **a,** Distribution of FOOOF fitting scores for hierarchy-consistency time series in the OMEGA dataset. **b,** Distribution of FOOOF fitting scores for hierarchy-consistency time series in the Cam-CAN dataset. In both independent cohorts, fitting scores were concentrated at high values, indicating that the spectral parameterization model reliably captured the spectral structure of hierarchy-consistency dynamics.

**Supplementary Fig. 2.**
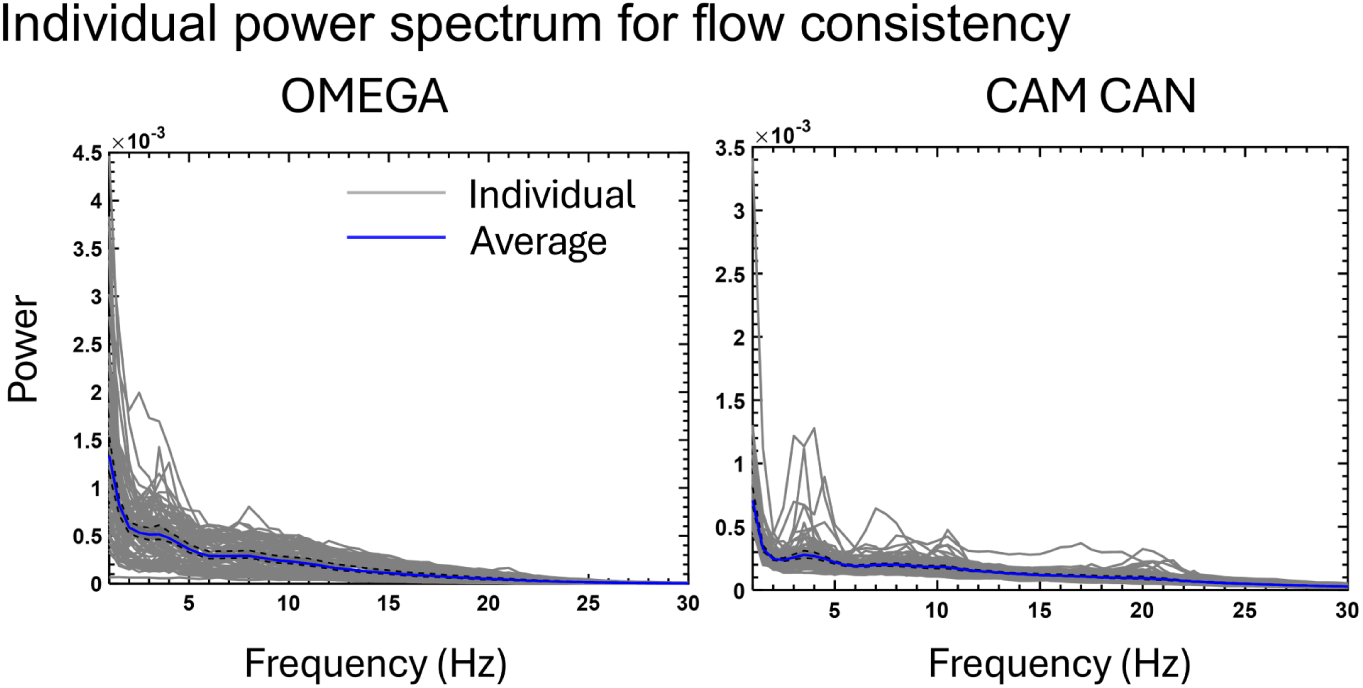
Individual power spectra of hierarchy-consistency dynamics. Power spectra of hierarchy-consistency time series are shown for the OMEGA and Cam-CAN datasets. Grey lines represent individual participants, and blue lines indicate the group-average spectrum. In both cohorts, hierarchy-consistency dynamics were dominated by low-frequency spectral structure, while also showing inter-individual variability in peak structure and spectral power.

**Supplementary Fig. 3.**
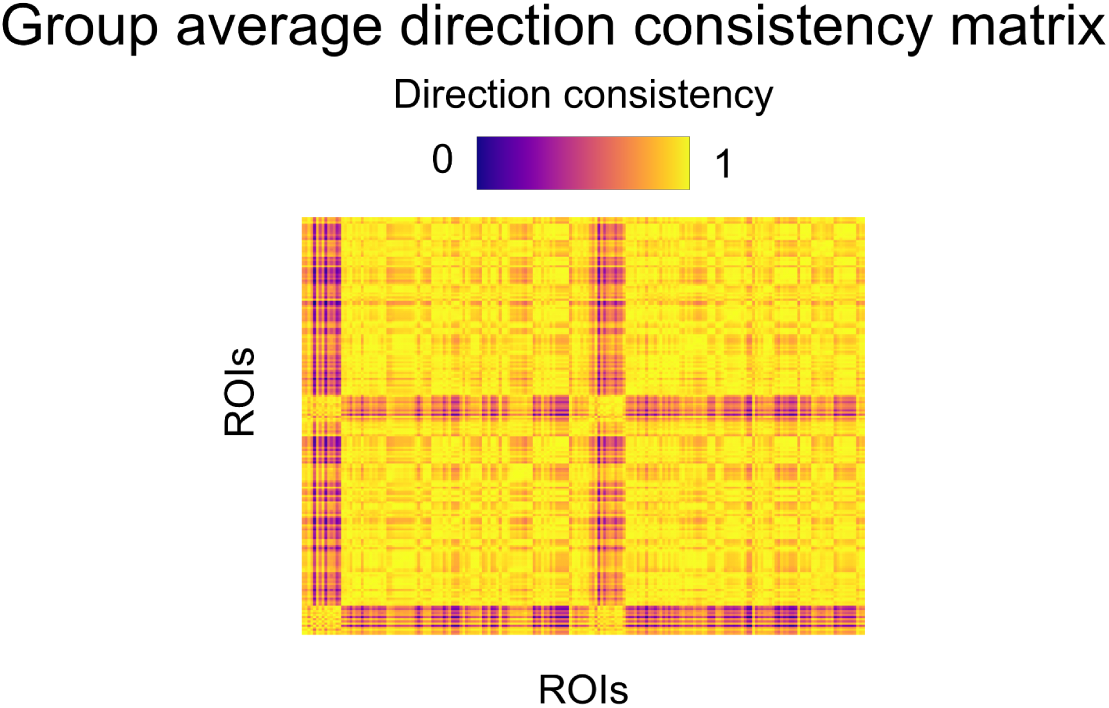
**Inter-regional synchronization of hierarchy-related cortical dynamics.**The matrix shows group-averaged phase-locking values between parcel-wise hierarchy-related temporal profiles. Rows and columns represent cortical ROIs, and colour indicates synchronization values ranging from 0 to 1. Higher values indicate stronger phase synchronization between regional hierarchy-related temporal profiles. The matrix shows a spatially structured organization of inter-regional synchronization in hierarchy-related cortical dynamics.

**Supplementary Fig. 4.**
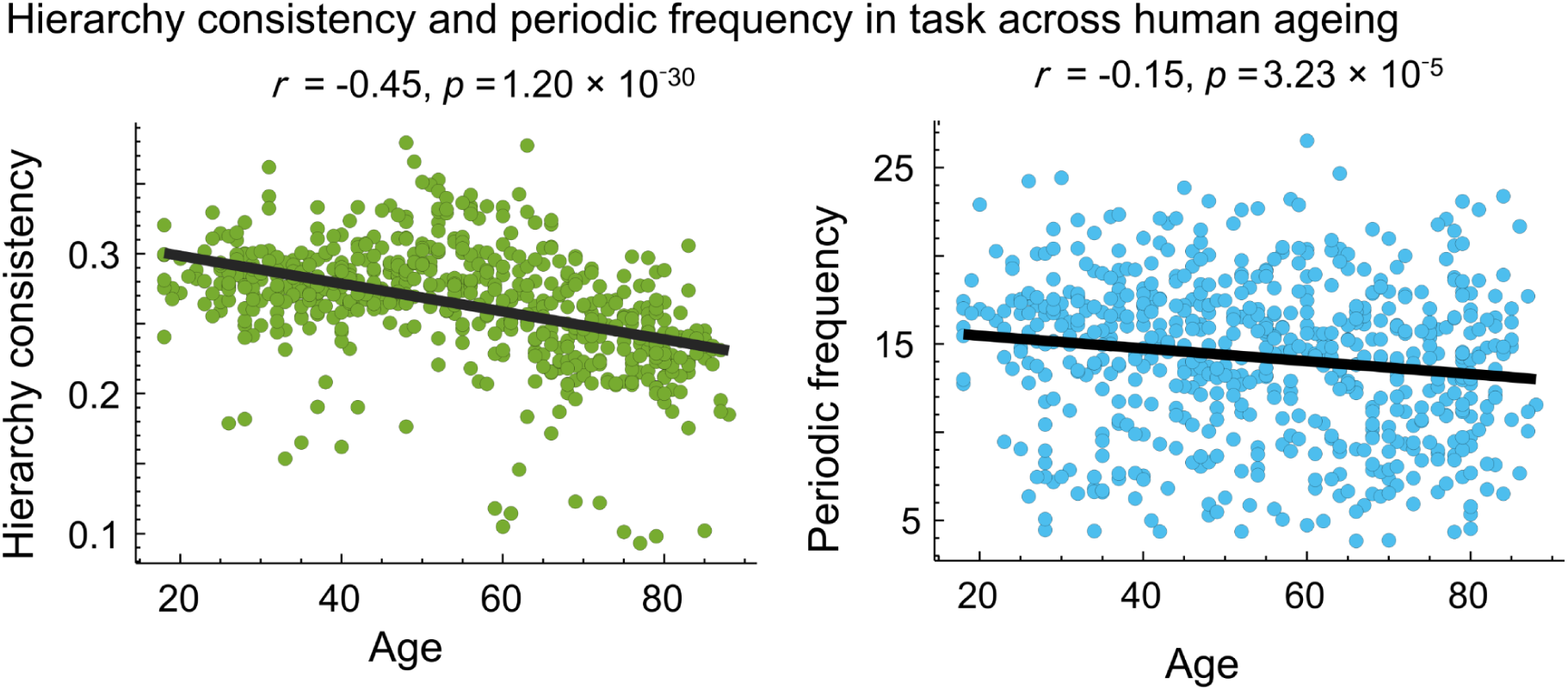
Hierarchy consistency and periodic frequency are associated with age. Age-related variation in task-derived hierarchy-consistency dynamics. Task hierarchy consistency decreased significantly with age (left; *r* = −0.45, *p* = 1.20 × 10⁻^30^), indicating weaker hierarchy-aligned propagation during task engagement in older adults. Task periodic frequency also decreased significantly with age (right; *r* = −0.15, *p* = 3.23 × 10⁻^5^), suggesting age-related slowing of hierarchy-related propagation dynamics during task. Each point represents one participant, and black lines indicate linear regression fits.

**Supplementary Fig. 5.**
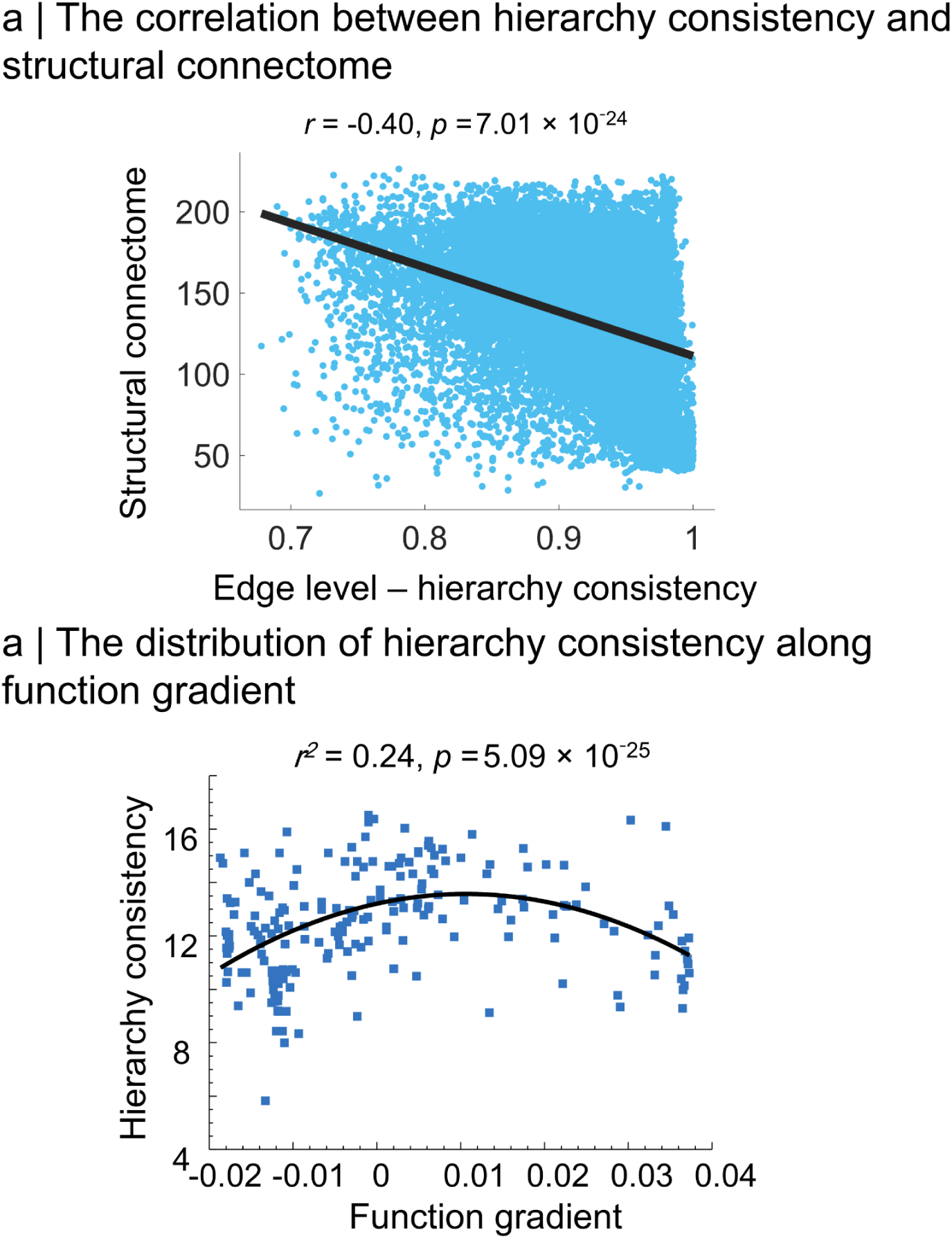
Structural and functional organization of hierarchy consistency. **a,** Relationship between edge-level hierarchy consistency and structural connectome measures. Edge-level hierarchy consistency was negatively associated with the structural connectome measure (*r = −0.40, p* = 7.01 × 10⁻^24^), indicating that edges with stronger consistency in propagation direction were preferentially associated with lower structural-connectome values. Each point represents a cortical edge, and the black line indicates the linear regression fit. **b,** Distribution of regional hierarchy consistency along the principal functional gradient. Hierarchy consistency showed a significant nonlinear association with the functional gradient (*r² = 0.24, p* = 5.09 × 10⁻^25^), indicating that hierarchy-consistent propagation is non-uniformly organized across the sensory-to-association axis. Each point represents a cortical region, and the black curve indicates the fitted nonlinear trend.

**Supplementary Fig. 6.**
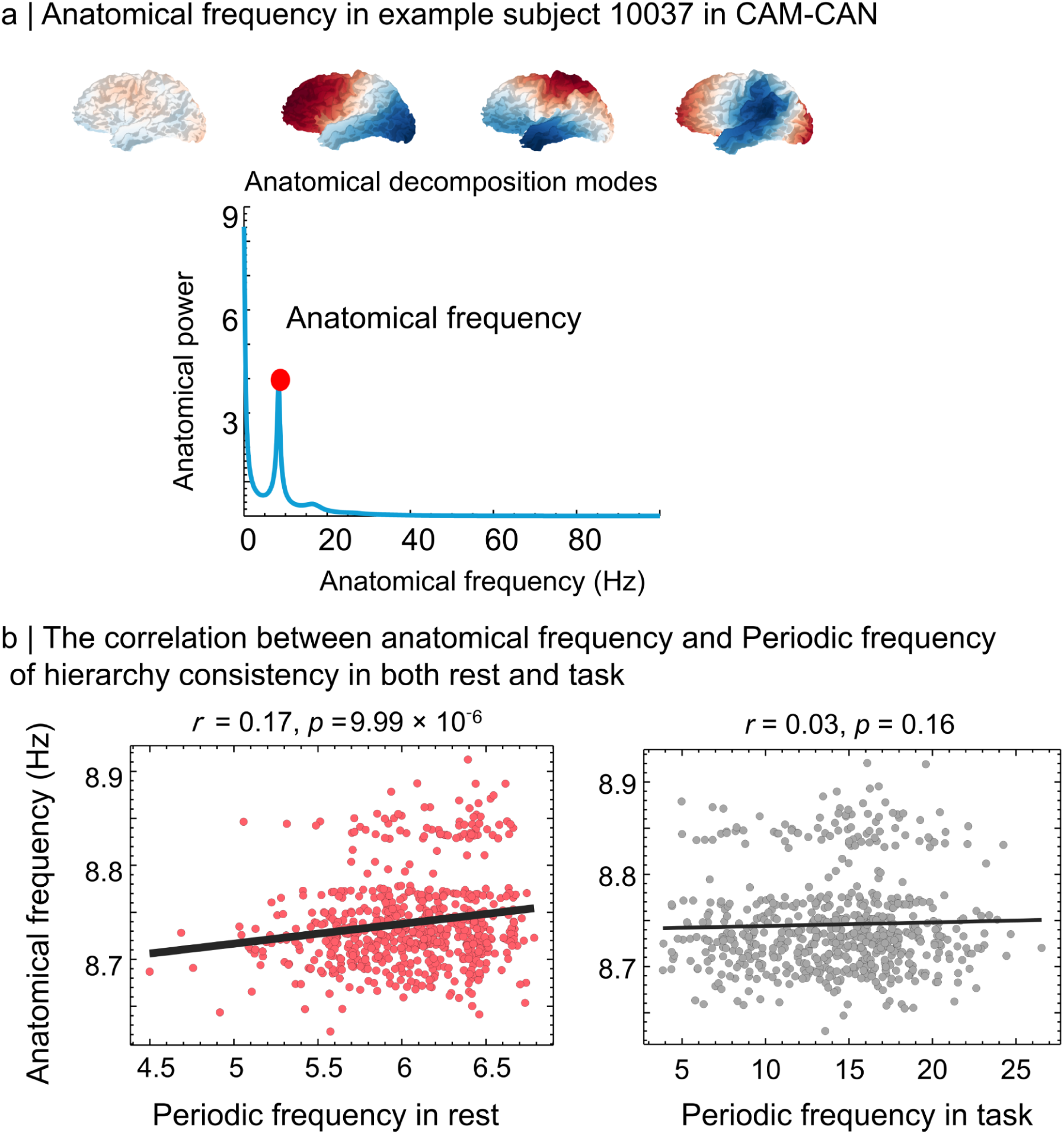
Anatomical frequency and its relationship with the periodic frequency of hierarchy consistency. **a,** Illustration of anatomical frequency estimation in an example Cam-CAN participant. Cortical anatomical decomposition modes were used to derive an anatomical power spectrum, and the dominant anatomical frequency was defined as the frequency corresponding to the largest spectral peak. The red dot marks the estimated anatomical frequency. **b,** Associations between anatomical frequency and the periodic frequency of hierarchy consistency during rest and task. Anatomical frequency showed a weak but significant positive association with the periodic frequency of hierarchy consistency during rest (*r* = 0.17, *p* = 9.99 × 10⁻^6^), whereas no significant association was observed during task (*r* = 0.03, *p* = 0.16). Each point represents one participant, and black lines indicate linear regression fits.

